# The MAASWERP study: An international, comparative case study on measuring biomechanics of the aged murine aorta

**DOI:** 10.1101/2024.09.17.613423

**Authors:** Cédric H.G. Neutel, Koen W.F. van der Laan, Callan D. Wesley, Dustin N. Krüger, Margarita G. Pencheva, Casper G. Schalkwijk, Guido R.Y. De Meyer, Wim Martinet, Tammo Delhaas, Koen D. Reesink, Alessandro Giudici, Pieter-Jan Guns, Bart Spronck

**Affiliations:** Laboratory of Physiopharmacology, University of Antwerp, Campus Drie Eiken, Antwerp, Belgium; Department of Biomedical Engineering, Cardiovascular Research Institute Maastricht (CARIM), Maastricht University, Maastricht, The Netherlands; Department of Internal Medicine, Cardiovascular Research Institute Maastricht (CARIM), Maastricht University, Maastricht, The Netherlands; GROW Institute for Oncology and Reproduction, Maastricht University, Maastricht, The Netherlands; Macquarie Medical School, Faculty of Medicine, Health and Human Sciences, Macquarie University, Sydney, NSW, Australia

**Author notes:** **<u>Corresponding author:</u>** Bart Spronck; Department of Biomedical Engineering, CARIM School for Cardiovascular Diseases, Maastricht University, Universiteitssingel 50,Room 3.356, 6229ER, Maastricht, The Netherlands. Authors contributed equally as first authors (Cédric H.G. Neutel, Koen W.F. van der Laan). Authors contributed equally as last authors (Pieter-Jan Guns, Bart Spronck).

**Keywords:** Aorta, Biomechanics, Methods

## Abstract

Arterial stiffening is a hallmark of vascular ageing, and unravelling its underlying mechanisms has become a central theme in the field of cardiovascular disease. While various techniques and experimental setups are accessible for investigating biomechanics of blood vessels both *in vivo* and *ex vivo*, comparing findings across diverse methodologies is challenging. In the present study, we aimed to compare arterial stiffness measurements of two distinct *ex vivo* setups for measuring aortic mechanics. First, we measured arterial stiffness in the aorta of adult (5 months) and aged (24 months) wild-type C57Bl/6J mice *in vivo*, after which *ex vivo* biomechanical evaluation was performed using the Rodent Oscillatory Tension Setup to study Arterial Compliance (ROTSAC; University of Antwerp, Belgium) and the DynamX setup (Maastricht University, The Netherlands). Measurements in both setups were conducted in parallel with matched protocols and identical buffers and chemicals. Overall, both methods revealed age-related increased stiffness, although parameters of aortic mechanics showed different numerical values, suggesting that results are not directly interchangeable between methods. Surprisingly, smooth muscle cell contraction had opposing effects between the setups. Indeed, smooth muscle cell contraction increased arterial stiffness in the ROTSAC but decreased stiffness in the DynamX. These opposing effects could be attributed to how the two setups differentially load the collagen fibres in the arterial wall, *ex vivo*. In conclusion, the observed differences between the two *ex vivo* setups highlight the necessity to report findings on (altered) aortic mechanics in the context of the used methodology.

## 2. Introduction

As the world’s population ages considerably, research on aging has become an ever-important field of science. Indeed, with an increase in the number of aged individuals, a parallel increase in the incidence of cardiovascular disease is expected, together with its associated economic burden [1, 2]. Arterial stiffening is a hallmark of cardiovascular aging and an important risk factor for multimorbidity in the aged population [3-5]. Therefore, investigating the mechanisms of aging-associated arterial stiffening is crucial for developing treatment options to reduce arterial stiffness and cardiovascular comorbidities in the elderly, which increases quality of life.

Despite its importance, at present, measuring “arterial stiffness” is (relatively) ambiguous. There are many methods available for measuring arterial stiffness in patients, yet clinically used parameters (mostly) provide general information without much mechanistic insight [6-8]. For instance, pulse wave velocity (PWV) is a commonly used metric for measuring arterial stiffness *in vivo*. Importantly, PWV is related to both material stiffness and vessel geometry, as well as their interaction [9]. Hence, an increased PWV could be the result of alterations in either parameter, or a combination of both. Furthermore, PWV is greatly defined by blood pressure, independent of changes in the underlying wall properties [10]. *Ex vivo* testing of aortic tissue allows for a much deeper investigation of the mechanisms of arterial stiffening, as it separates cellular and non-cellular contributions (i.e., matrix proteins) to arterial stiffness [11-14]. Different setups exist that measure arterial mechanics *ex vivo* [14, 15]. In brief, wire and pressure myograph systems are often employed to study both vascular reactivity as well as the biomechanical properties of arteries. However, comparing data generated by different setups, is difficult, as technical differences and differential output parameters (e.g., the representation of “stiffness”) hamper translatability among setups [14, 16-18]. Moreover, stiffness is often measured in static stretch conditions, which insufficiently reflect dynamic loading and unloading as observed *in vivo*.

In the present study, we compared the Rodent Oscillatory Tension Set-up to study Arterial Compliance (ROTSAC) at the University of Antwerp and the DynamX setup at Maastricht University, which are two distinct *ex vivo* setups for measuring murine arterial biomechanics in dynamic loading conditions [16, 17, 19]. In both setups we used fresh aortic tissue collected from the same mouse (of note, transport time from Belgium to The Netherlands needs to be taken into account here as well). Carefully synchronizing measurements at the two universities allowed for an optimal comparison with a minimum of confounding factors. Additionally, as both adult (i.e., 5-month-old) and aged (i.e., 24-month-old) mice were used, the present study also provides extensive insight into altered biomechanical properties of the aged murine aorta.

## 3. Methods

### 3.1 Animals

Adult (5-month-old, n=5) and aged (24-month-old, n=5) male C57Bl/6J mice (Charles River, France) were used for this study. All animals were housed in the animal facility of the University of Antwerp in standard cages with 12 h -12 h light-dark cycles and had free access to regular chow and tap water. All animal experiments were approved by the Ethical Committee of the University of Antwerp (ECD n° 2022/43) and were conducted in accordance with the EU Directive 2010/63/EU.

### 3.2 Chemicals

HEPES buffer (in mM) was prepared as follows: 143.3 NaCl, 4.7 KCl, 1.2 MgSO_4_.7H_2_O, 1.2 KH_2_PO_4_, 2.5 CaCl_2_, 5.5 Glucose, 15 HEPES. For studying vasoreactivity, phenylephrine, an α1-adrenoreceptor agonist, was used to elicit vascular smooth muscle cell (VSMC) contraction. L-N^G^-Nitroarginine methyl ester (L-NAME) was used to inhibit (endothelial) nitric oxide synthase activity and diethylamine nitric oxide (DEANO) was used to induce VSMC relaxation. Sodium nitroprusside (SNP) was used at Maastricht University to induce VSMC relaxation. Phenylephrine, L-NAME and DEANO were purchased from Sigma-Aldrich (Overijse, Belgium). SNP was purchased from Sigma-Aldrich (St. Louis, MO, USA).

### 3.3 Measurement of in vivo arterial stiffness

Ultrasound imaging was combined with invasive central blood pressure measurements to obtain the *in vivo* pressure–(inner) diameter relationship of the thoracic descending aorta. Blood pressure measurements were performed using a 1.4 F Mikro-Tip® mouse pressure catheter (Millar, SPR-671). Measurements were performed in the thoracic descending aorta. Ultrasound imaging was performed in anesthetized mice under 1.5–2.5% (*v*/*v*) isoflurane (Forene; Abbvie, Wavre, Belgium) using a high-frequency ultrasound system (Vevo F2 LAZR-X, VisualSonics, Toronto, Canada). Images were only acquired when heart rate and body temperature met the inclusion criteria, i.e., 500-600 beats/min and 36-38 °C, respectively. M-mode images were captured from the thoracic ascending aorta and thoracic descending aorta using a 24 MHz transducer to measure the inner diameter and wall thickness using Vevo LAB Software (Version 3.2.0, VisualSonics, Toronto, Canada).

### 3.4 *Measuring* ex vivo *aortic biomechanics*

*Ex vivo* biomechanical testing was performed using both a ROTSAC setup (at the University of Antwerp, Belgium) and a DynamX setup (at Maastricht University, The Netherlands) in parallel, as illustrated in **Figure 1**.

**Figure 1.**
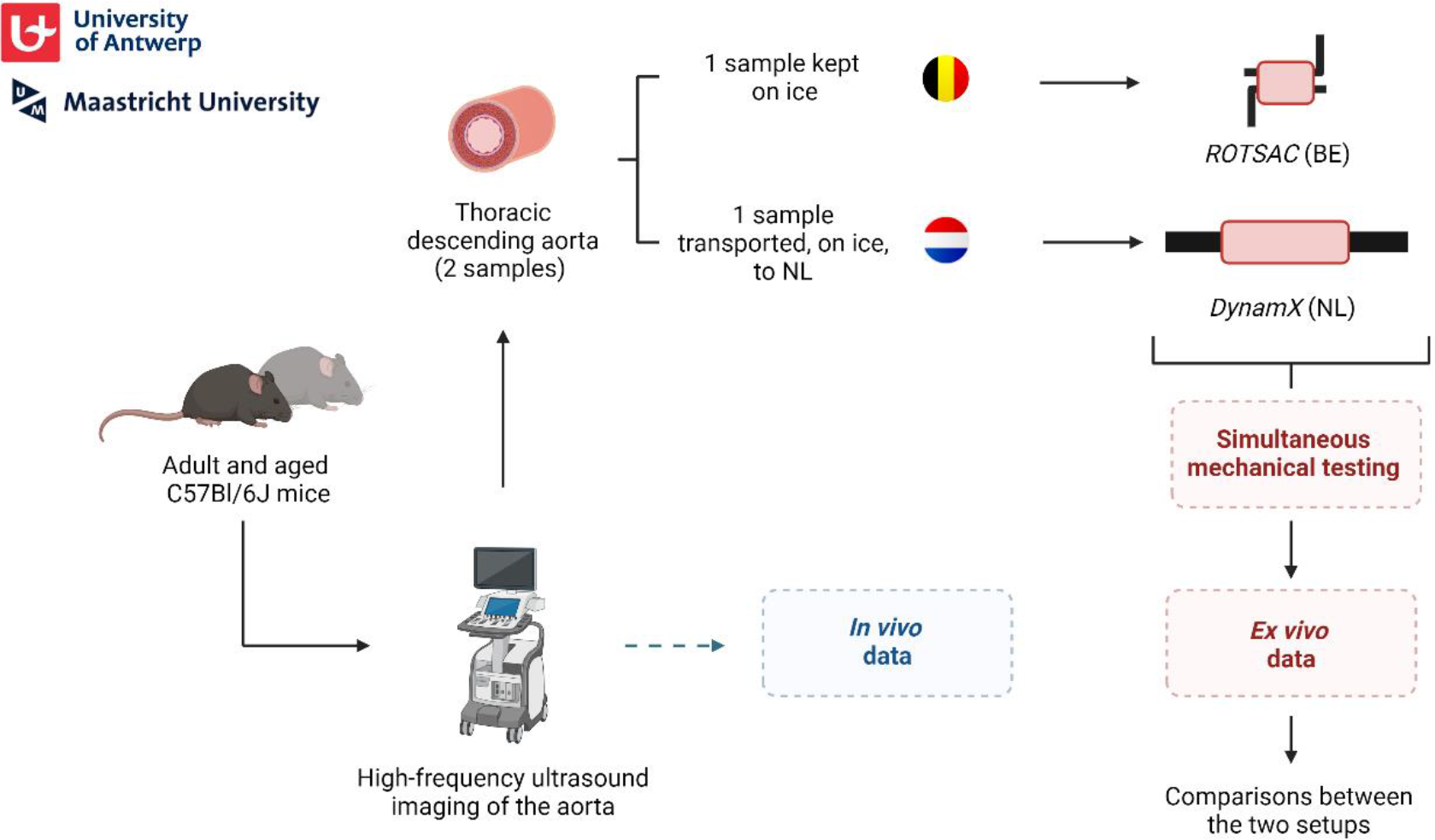
MAASWERP study protocol. *In vivo* arterial stiffness was determined using high-frequency ultrasound imaging combined with the measurement of central blood pressure using a pressure catheter. After *in vivo* imaging, two adjacent samples from the thoracic descending aorta were collected. One sample was kept on ice (in HEPES buffer) at the University of Antwerp (Belgium) whereas the other sample was transported on ice (in HEPES buffer) to Maastricht University (The Netherlands). Aortic samples were mounted in both *ex vivo* setups at the same time. Furthermore, mechanical testing in both setups was performed in parallel through constant communication between the two research groups to ensure data could directly be compared between both setups.

The ROTSAC is an in-house developed setup that allows for measuring aortic biomechanics and vascular reactivity [16]. In brief, 2 mm-long aortic segments were mounted between two parallel wire hooks in organ baths of 10 mL. Force and displacement of the upper hook were controlled and measured, respectively, with a force-and-length transducer. The segments were continuously oscillating between alternating preloads at a frequency of 10 Hz. Force and displacement were sampled at 400 Hz. Force was measured directly by the transducer. The equivalent inner diameter of the vessel segment (*d*_*i*_) at a given preload was derived from the inter-hook distance (*x*), being directly proportional to the inner circumference

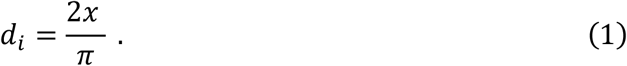

In this calculation, the thickness of the hooks between which the aortic segments were mounted was assumed to be negligible. Before each experiment, *x* and the vessel length (*l*) were determined statically at six different preloads (10, 20, 30, 40, 50, and 60 mN) using a camera and calibrated image software. To correct for the decrease in *l* with increased extrapolated diameter, the average length of the segment at each cycle (100 ms) was derived from the *d*_*i*_–*l* relationship using linear regression. The following relationship was used to calculate the transmural pressure from the distension force and vessel dimensions

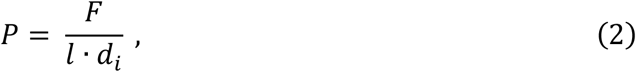

with *P* the pressure and *F* the force.

DynamX is a custom-built pressure myograph system capable of characterizing arterial biomechanics and vascular reactivity while mimicking the physiological axial stretch and pulsatile pressurization [17, 19]. Briefly, the aortic segment’s side branches were sutured closed (single knotted with nylon suture), after which the segment was mounted on two edged glass pipettes in the setup’s inner organ bath (double knotted with silk sutures).

Segments were stretched by motorized stages and pressurized using a combination of a pressure regulator and a pressure oscillator. Axial forces and luminal pressures were tracked using an *in situ* calibrated load cell connected to the distal glass pipette and an in-line distal pressure sensor, respectively, while a high-speed camera tracked the segment’s outer diameter. Any time delays between pressure experienced by the sample and measured by the sensor were eliminated by closing of the end of the pressure sensor and ensuring a short and rigid connection between the sample and sensor [17]. Before starting the experiments, segments were stretched to their *in vivo*-like length, defined as the length for which the change in measured axial force when pressurizing from 50 to 140 mmHg was minimal.

After mechanical testing in the DynamX system, three thin rings were cut from the distal end of the segment and submerged in a separate petri dish filled with HEPES buffer. Cross-section images were taken of these rings using a USB-camera (Dino-lite, Almere, The Netherlands) and stereomicroscope, from which segment wall thicknesses were determined. Assuming segment incompressibility, segment inner diameters were calculated according to

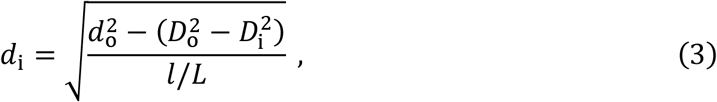

where *d*_o_ and *l* represent segment outer diameter and length, respectively, and *L*, 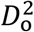, and 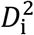 represent segment unloaded length, outer and inner diameter, respectively.

### 3.5 Study Protocol

The study protocol is illustrated in **Figure 1**. Ultrasound imaging and central blood pressure measurements were performed in the early morning. Mice were euthanized by perforating the diaphragm (performed under deep anaesthesia using 1.5–2.5% (v/v) isoflurane (Forene; Abbvie, Wavre, Belgium). Afterwards, the thoracic descending aorta was carefully removed from the mouse and stripped off all adherent tissue. Next, a 2 mm segment was cut distally and kept in HEPES buffer on ice in Antwerp, while the remainder larger part of the thoracic descending aorta was transported (in HEPES buffer on ice) to Maastricht University. Once in Maastricht, the aortic tissue was further processed and mounted in the biomechanical testing setup. Mounting of tissue segments was performed at the same time in the two setups. The experiments were also performed simultaneously through constant communication. In brief, aortic geometries and stiffness were assessed in basal, unstimulated conditions. Further, changes in diameter and stiffness were determined after inducing VSMC contraction using 2 μM PE (without and with addition of 300 μM L-NAME). Measurements were made with pressure oscillating between 70 and 110 mmHg, diastolic and systolic pressures respectively (“sinusoidal” loading waveform). Lastly, vasorelaxation was induced using 2 μM DEANO or 10 μM SNP in ROTSAC and DynamX, respectively. After inducing VSMC relaxation, the pressure–stiffness relationship was studied by incrementally increasing the mean pressure in steps of 20 mmHg, keeping a constant dynamic pressure pulse of 40 mmHg amplitude. Phenylephrine and L-NAME stocks were prepared in Antwerp and were transported to Maastricht. Therefore, not only the same aortic tissue was used, but also buffers and stock solutions of vasoactive compounds were identical (with exception of DEANO, used at University of Antwerp, and SNP, used at Maastricht University, to induce vasorelaxation). This protocol was repeated for every mouse in this study.

### 3.6 Calculation of stiffness variables

The Peterson’s modulus of elasticity (*E*_P_) was chosen as a quantifier of *in vivo* and *ex vivo* aortic stiffness and was calculated as

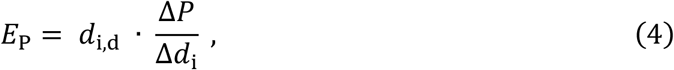

with *d*_i,d_ the inner diastolic diameter, Δ*d*_i_ the distension and Δ*P* the pulse pressure. Additionally, aortic compliance (*C*) was determined and was calculated as:

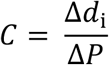

### 3.7 Statistics

Statistical analyses were performed in GraphPad Prism 10. All data are presented as mean ± standard error of the mean with *n* representing the number of mice. Statistical tests are specified in the figure legends. Differences were considered significant when *p*<0.05.

## 4. Results

### 4.1 Ageing increases arterial stiffness *in vivo* in mice

Vascular parameters (on the thoracic descending aorta) were determined for both adult (5-month-old) and aged (24-month-old) mice and are summarized in **Table 1**. While no differences were observed in aortic diameter and central blood pressure, the aortic compliance and Peterson’s modulus of elasticity were significantly decreased (*p*=0.015) and increased (*p*=0.017), respectively, in aged compared to adult mice, indicative for increased structural arterial stiffness.

**Table 1.**
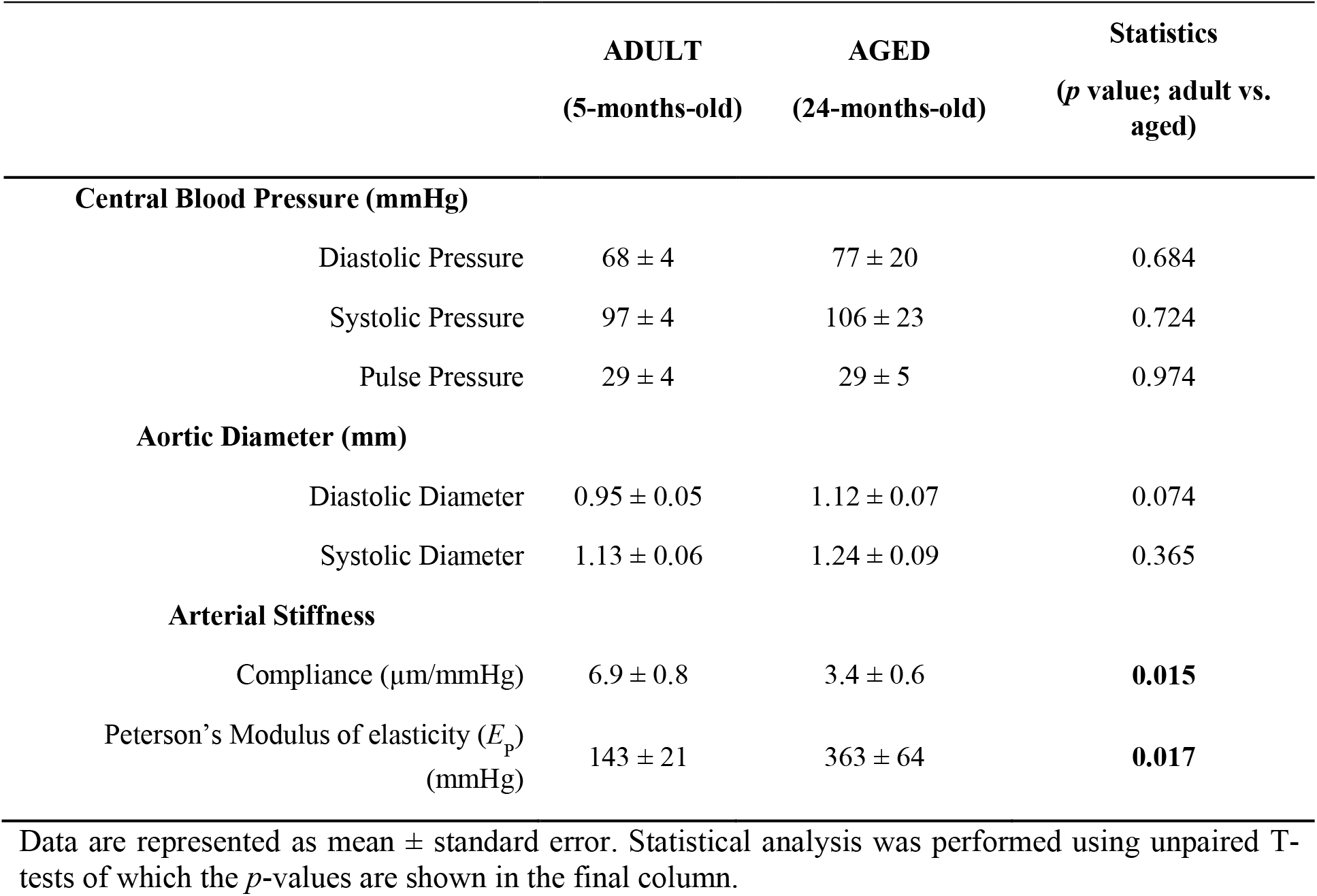
*In vivo* vascular characteristics of adult and aged C57Bl6/J mice.

|  | ADULT<br>(5-months-old) | AGED<br>(24-months-old) | Statistics<br>(*p* value; adult vs.<br>aged) |
| --- | --- | --- | --- |
| **Central Blood Pressure (mmHg)** |  |  |  |
| Diastolic Pressure | 68 ± 4 | 77 ± 20 | 0.684 |
| Systolic Pressure | 97 ± 4 | 106 ± 23 | 0.724 |
| Pulse Pressure | 29 ± 4 | 29 ± 5 | 0.974 |
| **Aortic Diameter (mm)** |  |  |  |
| Diastolic Diameter | 0.95 ± 0.05 | 1.12 ± 0.07 | 0.074 |
| Systolic Diameter | 1.13 ± 0.06 | 1.24 ± 0.09 | 0.365 |
| **Arterial Stiffness** |  |  |  |
| Compliance (µm/mmHg) | 6.9 ± 0.8 | 3.4 ± 0.6 | **0.015** |
| Peterson's Modulus of elasticity ($E_p$)<br>(mmHg) | 143 ± 21 | 363 ± 64 | **0.017** |

Data are represented as mean ± standard error. Statistical analysis was performed using unpaired T-tests of which the *p*-values are shown in the final column.

### 4.2 Comparison between the ROTSAC and DynamX setups in measuring vascular ageing *ex vivo*

Aortic samples from adult and aged C57Bl6/J mice were measured in both setups, in parallel, in unstimulated conditions. Both (internal) diameters and stiffness of the aortic samples were measured and compared to each other (**Figure 2**). First, whereas diastolic diameters significantly increased with ageing (*p*<0.01), multiple comparison analysis revealed that the increase was significant in the ROTSAC (*p*<0.01) but not in the DynamX (**Figure 2A**). Furthermore, internal diameters of aortic samples were in general significantly larger in the DynamX compared to the ROTSAC (*p*<0.0001). While significant differences (p<0.01) in estimated stiffness were observed between the two setups, the following consistent observations could be made: arterial stiffness (expressed as *E*_P_) was also increased with ageing (**Figure 2B**). However, multiple comparison analysis revealed that the increase was significant in the DynamX (*p*<0.0001) but not in the ROTSAC.

**Figure 2.**
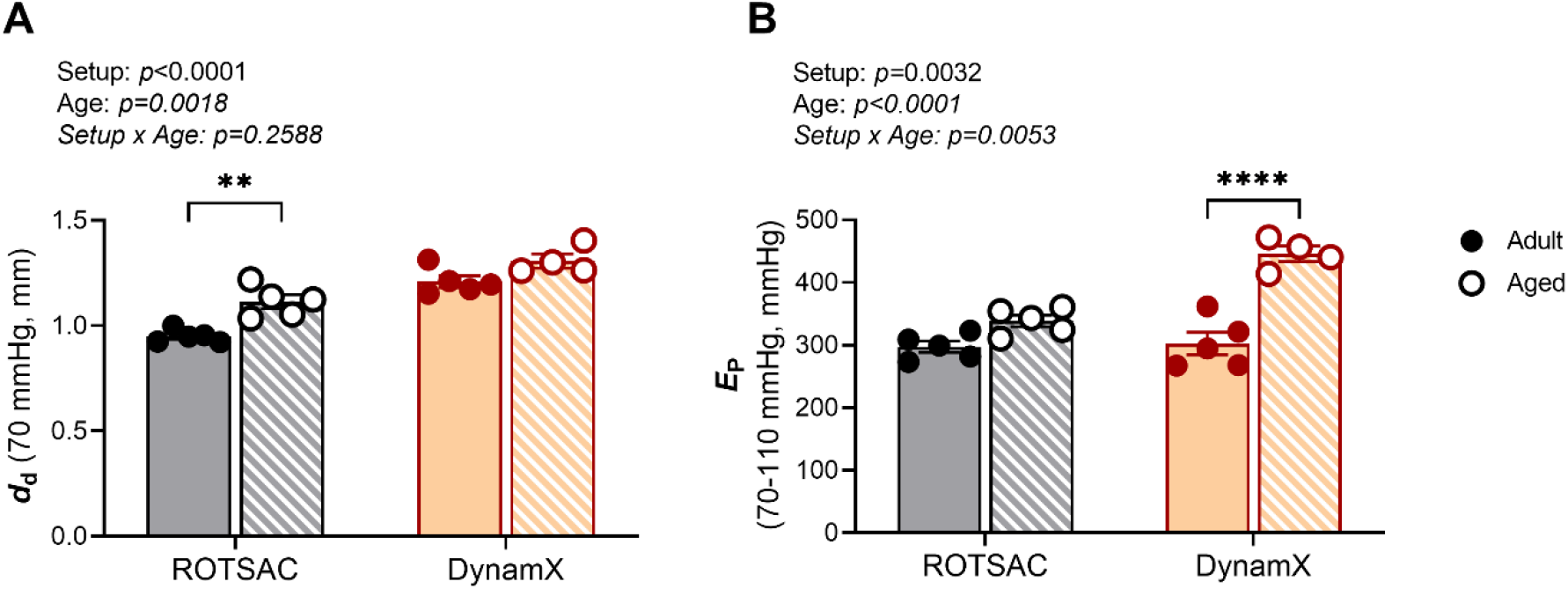
Comparison between ROTSAC and the DynamX setups in measuring vascular ageing *ex vivo*. (A) Diastolic diameter measured at 70 mmHg. (B) Aortic stiffness, measured as Peterson’s modulus of elasticity (E_P_). Statistical analysis was performed using a two-way analysis of variance (ANOVA) with a Tukey post hoc test for multiple comparisons. *n*=4–5. \*\**p*<0.01, ****p<0.0001. *d*_d_ = diastolic diameter; *E*_P_ = Peterson’s modulus of elasticity.

### 4.3 The *ex vivo* pressure–stiffness relationship exhibits distinct differences between the two experimental setups

Next, the pressure–stiffness relationship was evaluated after inducing vasorelaxation, using 2 μM DEANO or 10 μM SNP in the ROTSAC and DynamX, respectively (**Figure 3**). By inducing vasorelaxation, passive mechanical behaviour was measured. Incrementally increasing mean pressure (at a constant pulse pressure of 40 mmHg) increased the aortic (internal) diameter and tissue stiffness. An ageing effect was observed on both *ex vivo* aortic diameter (p<0.01) and stiffness (p<0001) in the ROTSAC (**Figure 3A, B**). However, no ageing effect was observed on either diameter or stiffness in the DynamX (**Figure 3C, D**). Alternatively, a significant interaction (*p*<0.0001) between the pressure and diameter parameters was observed in the DynamX. Indeed, at 50 mmHg, the diameters of aortic samples of aged mice were significantly (*p*<0.01) larger than those originating from adult mice.

**Figure 3.**
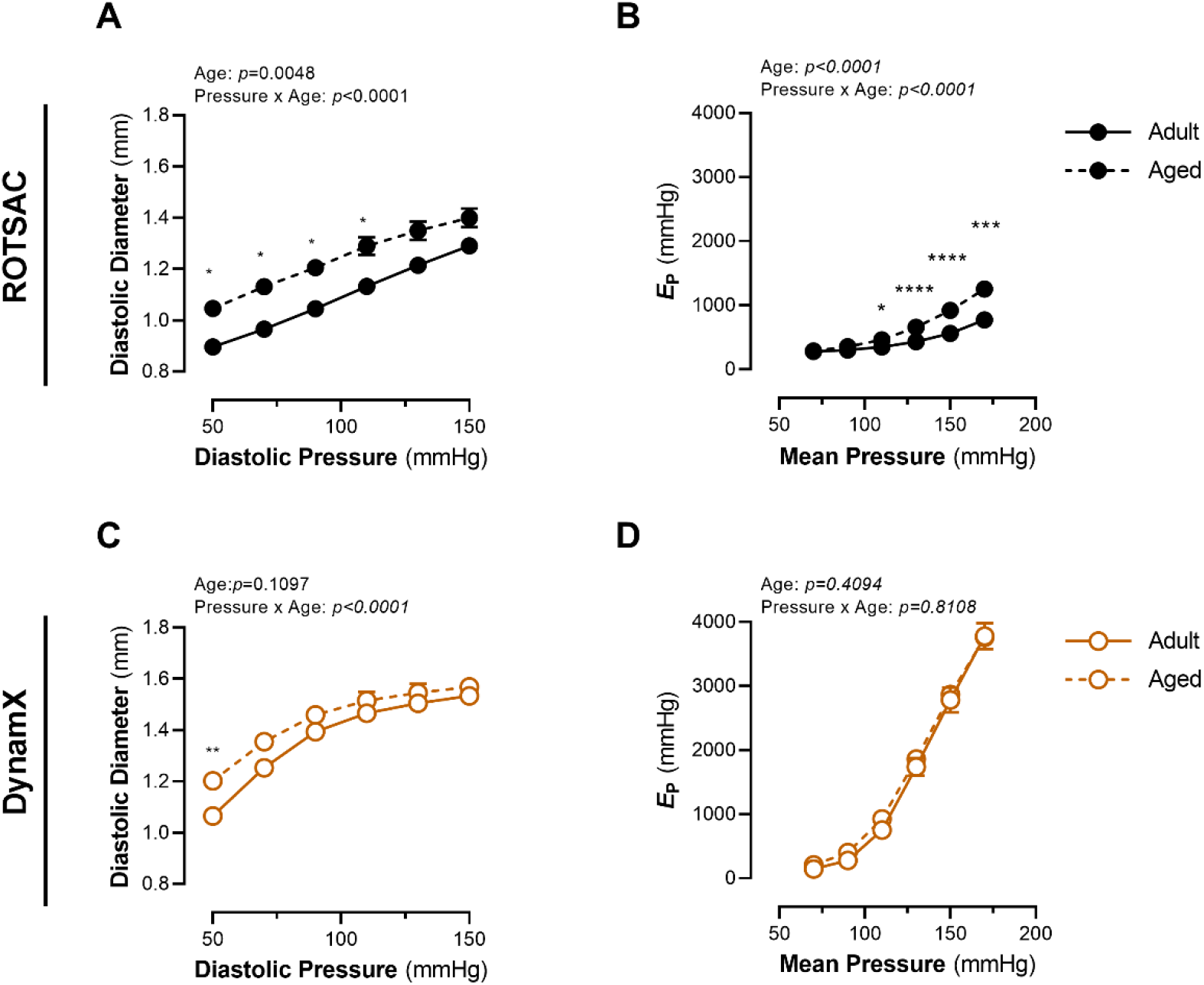
Passive aortic biomechanics in the ROTSAC and in the DynamX setup. Aortic samples from both adult and aged C57Bl6/J mice were subjected to incrementally increasing systolic and diastolic pressures (at a constant pulse pressure of 40 mmHg) in both the ROTSAC and the DynamX, in parallel. Increasing “pressure” in the ROTSAC increases the (A) diameter and (B) stiffness of the aortic samples. Similarly, increasing pressure in the DynamX increases both (C) the diameter and (D) stiffness of the aortic samples. *n*=4–5. Statistical analysis was performed using a two-way ANOVA with a Sidak post hoc test for multiple comparisons. \*\**p*<0.01, \*\*\**p*<0.001, ****p<0.0001. *E*_P_ = Peterson’s modulus of elasticity.

### 4.4 *Ex vivo* vascular smooth muscle cell contraction modulates aortic tissue biomechanics differently in the two setups

VSMC contraction was induced with 2 μM phenylephrine (PE) under cyclic stretch. Afterwards, the change in (internal) diastolic diameter as well as changes in aortic stiffness, determined through changes in the Peterson’s modulus, were measured between 70 and 110 mmHg diastolic and systolic pressure, respectively (**Figure 4**). In the ROTSAC, VSMC contraction decreased internal diameter and increased arterial stiffness (**Figure 4A, B**). Moreso, an ageing effect was observed on the effect of VSMC on internal diameter (*p*<0.01) as well as on stiffness (*p*<0.05). Surprisingly, VSMC contraction resulted in an opposite stiffness response in the DynamX. Here, VSMC contraction decreased internal diameter, but also arterial stiffness (**Figure 4C, D**). A significant ageing effect (regarding the diameter change (p<0.05) and stiffness (p<0.001)) was observed in the DynamX as well.

**Figure 4.**
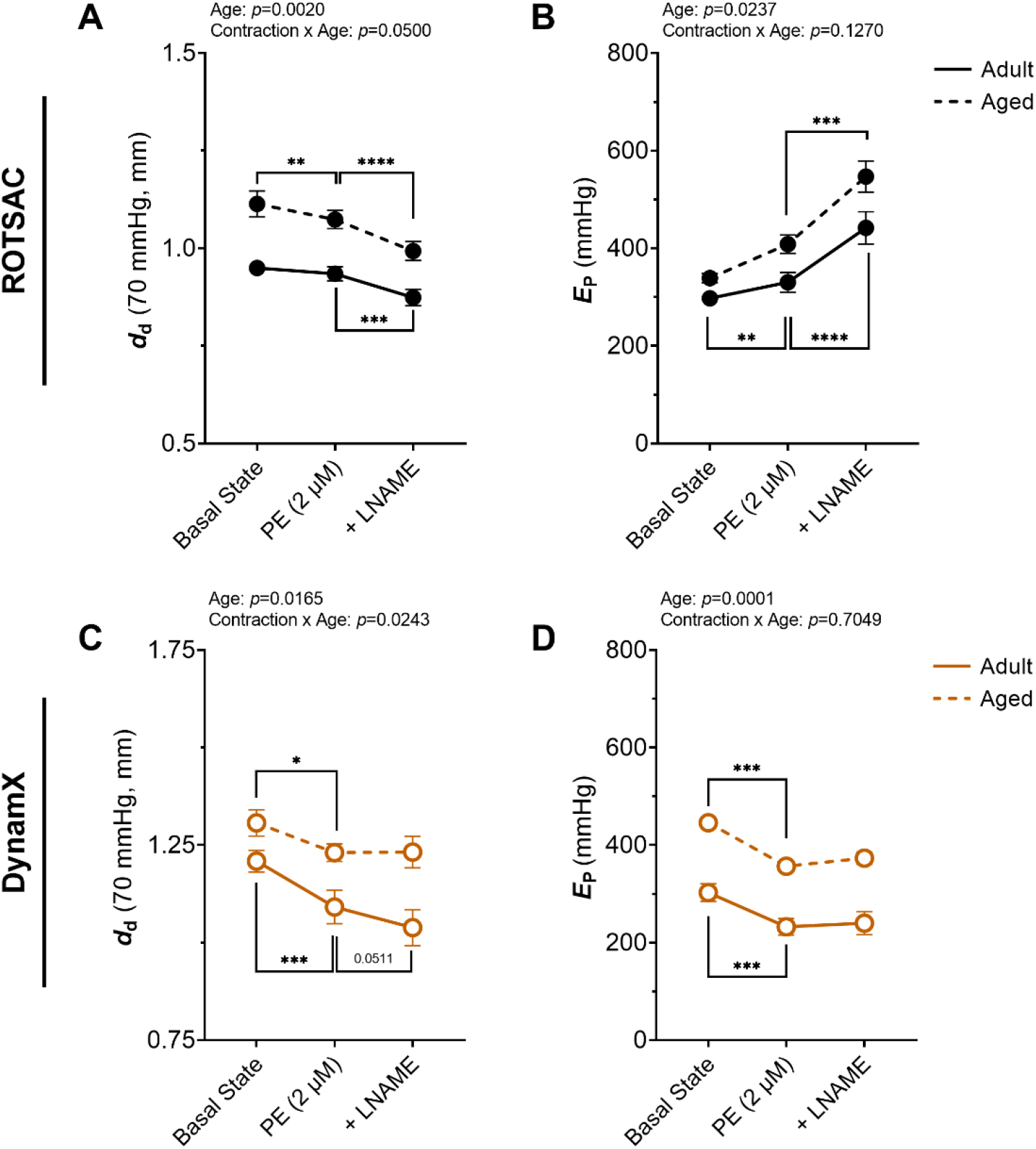
Differential effect of VSMC contraction on *ex vivo* arterial biomechanics between ROTSAC and DynamX. To investigate how VSMC can actively modulate aortic mechanics, VSMC contraction was elicited in both setups. First, a contraction was induced with 2 μM PE, after which 300 μM LNAME was added to reach maximum contraction. (A, B) In the ROTSAC, VSMC contraction decreased diameter, but gradually increased stiffness. (C, D) Also in the DynamX, the diameter of the aortic samples decreased after VSMC contraction, but, in stark contrast with the ROTSAC, VSMC contraction decreased arterial stiffness. Statistical analysis was performed using a paired two-way ANOVA with a Holm-Sidak post hoc test for multiple comparisons. *n*=4–5. *p<0.05, \*\**p*<0.01, \*\*\**p*<0.001, \*\*\*\**p*<0.0001. *d*_d_ = diastolic inner diameter; *E*_P_ = Peterson’s Modulus; PE = phenylephrine; L-NAME = L-N^G^-Nitro arginine methyl ester.

### 4.5 A technical correction for the curvature of the hooks in the ROTSAC only partially explains the numerical differences between the two setups

In an attempt to explain the dissonant results between the setups, a correction factor was applied to the ROTSAC which accounts for the hook size (**Supplementary Data 1**). This correction resulted in minor increases in both the diameter and stiffness values in the ROTSAC (**Supplementary Table 1**). Nevertheless, the corrected ROTSAC values still significantly differed from those obtained using the DynamX setup.

## 5. Discussion

The primary aim of this study was to evaluate the differences between distinct setups for measuring *ex vivo* murine aortic stiffness. To this end, we performed a study with the Antwerp (BE)-based ROTSAC and the Maastricht (NL)-based DynamX setup in measuring arterial mechanics of (murine) aortic samples. In both setups, aortic samples undergo high-frequency cyclic stretching, which significantly enhances the translation to *in vivo* conditions, presenting a major advantage over conventional static loading methods. However, whereas aortic samples are stretched uniaxially in the ROTSAC, they are stretched bi-axially in the DynamX. One can imagine that such a different mode of stretching has implications on its biomechanical properties. However, how much do such measurements really differ? In the next sections, the major differences are discussed, as well as how vascular ageing presented itself in both setups. Indeed, as samples from adult and aged mice were collected and measured, both setups were additionally benchmarked in their capacity to detect vascular ageing.

### 5.1 DynamX vs. ROTSAC: a matter of collagen loading?

In the ROTSAC, aortic segments are cyclically stretched in the circumferential direction in the absence of axial pre-stretch. In essence, the ROTSAC is an adaptation of wire myography. However, it must be noted that in ROTSAC the sample length shrinks in response to circumferential loading. Moreover, during loading, the sample is shorter than its unloaded length, implying that the sample is axially compressed (as opposed to the *in vivo* situation, where an artery is continuously axially extended). This subphysiological sample length affects the sample’s mechanical properties and results in the underestimation of the load on the different arterial wall layers [13]. In contrast, the DynamX setup is more similar to pulsatile pressure myographs. In DynamX, aortic segments are distended by the intraluminal pressure in the presence of an axial pre-stretch similar in magnitude to that of the artery *in vivo*.

Overall, measured/estimated inner diastolic diameters and tissue stiffness were greater in the DynamX compared to the ROTSAC. In addition, the pressure–stiffness relationship was much steeper in DynamX than in ROTSAC. This difference could be caused by the difference in axial loading between the setups. Indeed, unlike in the ROTSAC, axial length is kept constant at a value >1 in the DynamX (i.e., with the vessel extended compared to its unloaded state). Axial (pre-)stretch is known to modulate aortic stiffness and vascular contractility [17, 20]. Increasing (axial) stretch enhances arterial stiffness by reorganizing collagen fibres towards the direction of the load, reducing their waviness and therefore increasing total tissue stiffness [21-23]. In DynamX, collagen fibres are already substantially loaded at physiological pressures due to the axial prestretch. Therefore, increasing pressure leads to further loading on already (partially) stretched collagen fibres, resulting in large increases in *E*_P_. Conversely, in ROTSAC, collagen fibres are presumably less stretched (especially at lower pressures) due to the absence of axial pre-stretch. Therefore, increasing pressure in ROTSAC leads to relatively modest changes in *E*_P_ (compared to DynamX) as the collagen fibres slowly switch from a “wavy” configuration to a “stretched” shape with increasing pressure. This may explain why the pressure–stiffness relationship is different in the two setups.

The effect of the axial stretch on collagen fibre recruitment could also explain the contrasting results regarding the effect of VSMC contraction and aortic stiffness. Contraction of VSMCs in the aorta shifts the load from the extracellular matrix (ECM) to the VSMCs [24]. In DynamX, because of the superimposed effects of axial and circumferential stretches, the stiffness of the ECM at physiological pressures is determined by both elastin and stretched collagen fibres. Conversely, in ROTSAC, elastin is likely the main load-bearing component at these pressures. In DynamX, VSMC contraction leads to smaller diameters, thereby unloading the (stiff) collagen fibres and resulting in a decrease in stiffness. Conversely, in ROTSAC, VSMC contraction increases stiffness as load is being transferred from the distensible elastin fibres to the stiffer, contracted VSMCs. This hypothesis is further corroborated by previous studies of our group, which have shown that in ROTSAC, at higher pressures (i.e., greater collagen loading) VSMC contraction decreases aortic stiffness [24], while at lower pressures VSMCs contraction increases aortic stiffness.

### 5.2 Both setups detect ageing-induced changes in arterial properties, but with different sensitivities

A second question was whether both setups detect similar changes between aortic samples from adult and aged mice. This is important, as researchers who investigate *ex vivo* aortic biomechanics in disease models and/or try to identify whether a pharmacological therapy to reduce arterial stiffening was effective, would often only use one *ex vivo* method. First, we re-established that natural ageing induces arterial stiffness *in vivo*, which was a prerequisite for this study [11, 25]. Second, we have demonstrated that the DynamX and ROTSAC have slightly different sensitivity concerning vascular ageing. Indeed, while aortic samples from aged mice exhibited increased stiffness under basal but not passive conditions in the DynamX, the ROTSAC produced opposing results. In the ROTSAC, the stiffness of “aged” aortic samples was higher under passive but not basal conditions. Whereas both setups were able to identify increased stiffness with ageing, they did so in a different way. This could be explained by the different type of stretches and conditions the aortic samples are subjected to in the two setups. Hence, interpreting the data requires careful consideration and should be contextualized based on the methodology used. Therefore, this study highlights the need for reporting results in the context of the method employed. Indeed, dissonant results in measuring aortic biomechanics, obtained between different setups, do not necessarily mean “contradictory results” as they could present the same finding observed through a different methodological angle.

## 6. Conclusion

This study compared *ex vivo* measurements of murine aortic biomechanics between two different setups using tissue samples originating from the same aorta. The stiffness of the aortic tissue, expressed as Peterson’s modulus, differed between the setups in both basal, unstimulated conditions and passive conditions. Remarkably, VSMC contraction elicited opposing effects on the properties of the aortic tissue, increasing stiffness in the ROTSAC while decreasing stiffness in the DynamX. Overall, this study provided critical insights into how different experimental setups can influence the interpretation of aortic biomechanics, emphasizing the need for careful consideration and contextualization of results based on the methodology used.

## 9. Data availability statement

The data that support the findings of this study are available from the corresponding author, upon reasonable request.

## 10. CRediT authorship contribution statement

**Cédric H.G. Neutel:** Conceptualization, Methodology, Investigation, Formal analysis, Data Curation, Writing - Original Draft. **Koen W.F. van der Laan**: Conceptualization; Methodology, Software, Investigation, Formal analysis, Writing - Review & Editing. **Callan D. Wesley:** Investigation, Formal analysis, Writing - Review & Editing. **Dustin N. Krüger**: Investigation, Formal analysis, Writing - Review & Editing. **Margarita G. Pencheva**: Investigation, Formal analysis, Writing - Review & Editing. **Casper G. Schalkwijk**: Formal analysis, Writing - Review & Editing. **Guido R.Y. De Meyer**: Formal analysis, Writing - Review & Editing. **Wim Martinet**: Formal analysis, Writing - Review & Editing. **Tammo Delhaas**: Formal analysis, Writing - Review & Editing. **Koen D. Reesink**: Formal analysis, Writing - Review & Editing. **Alessandro Giudici**: Methodology, Software, Investigation, Formal analysis, Writing - Review & Editing. **Pieter-Jan Guns**: Conceptualization, Methodology, Writing - Review & Editing, Project administration, Funding acquisition. **Bart Spronck**: Conceptualization, Methodology, Writing - Review & Editing, Project administration, Funding acquisition

## SUPPLEMENTARY MATERIAL

### Supplementary Data 1

To correct for hook size in the ROTSAC, the formula to calculate the internal diameter was adjusted:

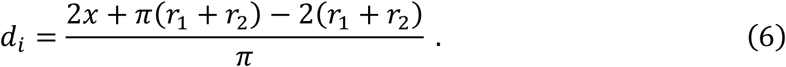

Where *d*_*i*_ and *x* denote the inner diameter of the vessel segment and the distance between the hooks, respectively, and *r*_1_ and *r*_2_ indicate the radii of the two hooks.

### Supplementary Table 1

**Supplementary Table 1.**
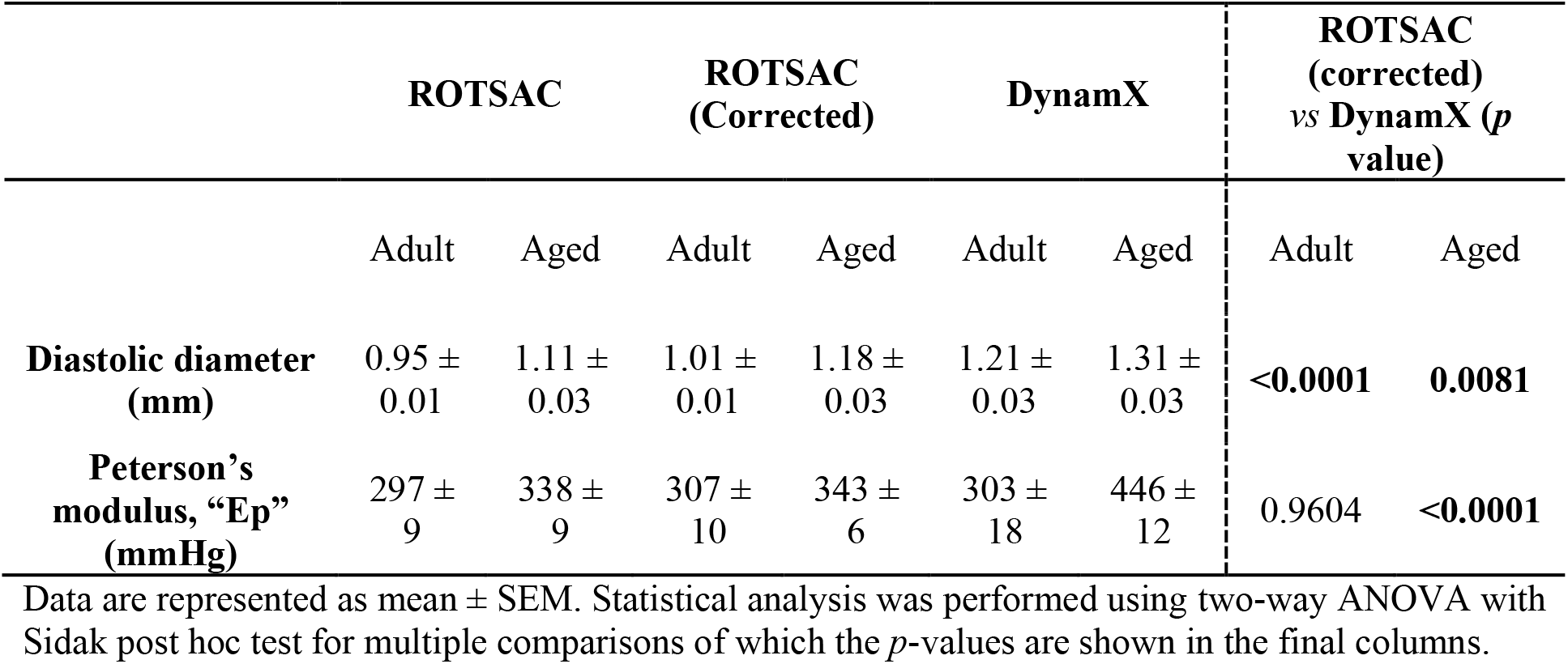
Correcting for hook size in the ROTSAC.

|  | ROTSAC |  | ROTSAC<br>(Corrected) |  | DynamX |  | ROTSAC<br>(corrected)<br>vs DynamX (*p*<br>value) |  |
| --- | --- | --- | --- | --- | --- | --- | --- | --- |
|  | Adult | Aged | Adult | Aged | Adult | Aged | Adult | Aged |
| **Diastolic diameter**<br>**(mm)** | 0.95 ±<br>0.01 | 1.11 ±<br>0.03 | 1.01 ±<br>0.01 | 1.18 ±<br>0.03 | 1.21 ±<br>0.03 | 1.31 ±<br>0.03 | **<0.0001** | **0.0081** |
| **Peterson's**<br>**modulus, "Ep"**<br>**(mmHg)** | 297 ±<br>9 | 338 ±<br>9 | 307 ±<br>10 | 343 ±<br>6 | 303 ±<br>18 | 446 ±<br>12 | 0.9604 | **<0.0001** |

Data are represented as mean ± SEM. Statistical analysis was performed using two-way ANOVA with Sidak post hoc test for multiple comparisons of which the *p*-values are shown in the final columns.

## References

1. National Institute On Aging. Heart Health and Aging. 2018 1 Jun 2018 [cited 2023; Available from: https://www.nia.nih.gov/health/heart-health-and-aging#:~:text=The%20most%20common%20aging%20change,more%20common%20as%20we%20age.

2. North, B.J. and D.A. Sinclair, The intersection between aging and cardiovascular disease. Circ Res, 2012. 110(8): p. 1097–108.

3. Mitchell, G.F., Arterial Stiffness in Aging: Does It Have a Place in Clinical Practice? Hypertension, 2021. 77(3): p. 768–780.

4. Mitchell, G.F., et al., Arterial stiffness and cardiovascular events: the Framingham Heart Study. Circulation, 2010. 121(4): p. 505–11.

5. Lee, H.Y. and B.H. Oh, Aging and arterial stiffness. Circ J, 2010. 74(11): p. 2257–62.

6. Segers, P., E.R. Rietzschel, and J.A. Chirinos, How to Measure Arterial Stiffness in Humans. Arterioscler Thromb Vasc Biol, 2020. 40(5): p. 1034–1043.

7. Cheng, H.-M. and C.-H. Chen, Measuring arterial stiffness in clinical practice: Moving one step forward. The Journal of Clinical Hypertension, 2020. 22(10): p. 1824–1826.

8. Budoff, M.J., et al., Clinical Applications Measuring Arterial Stiffness: An Expert Consensus for the Application of Cardio-Ankle Vascular Index. Am J Hypertens, 2022. 35(5): p. 441–453.

9. Fabian, V., et al., Noninvasive Assessment of Aortic Pulse Wave Velocity by the Brachial Occlusion-Cuff Technique: Comparative Study. Sensors (Basel), 2019. 19(16).

10. Spronck, B., et al., Pressure-dependence of arterial stiffness: potential clinical implications. Journal of Hypertension, 2015. 33(2): p. 330–338.

11. De Moudt, S., et al., Progressive aortic stiffness in aging C57Bl/6 mice displays altered contractile behaviour and extracellular matrix changes. Commun Biol, 2022. 5(1): p. 605.

12. Sulejmani, F., et al., Biomechanical properties of the thoracic aorta in Marfan patients. Annals of Cardiothoracic Surgery, 2017. 6(6): p. 610–624.

13. Giudici, A., et al., Tri-layered constitutive modelling unveils functional differences between the pig ascending and lower thoracic aorta. Journal of the Mechanical Behavior of Biomedical Materials, 2023. 141: p. 105752.

14. Butlin, M., et al., Measuring Arterial Stiffness in Animal Experimental Studies. Arterioscler Thromb Vasc Biol, 2020. 40(5): p. 1068–1077.

15. Giudici, A., I.B. Wilkinson, and A.W. Khir, Review of the Techniques Used for Investigating the Role Elastin and Collagen Play in Arterial Wall Mechanics. IEEE Rev Biomed Eng, 2021. 14: p. 256–269.

16. Leloup, A.J., et al., A novel set-up for the ex vivo analysis of mechanical properties of mouse aortic segments stretched at physiological pressure and frequency. J Physiol, 2016. 594(21): p. 6105–6115.

17. van der Bruggen, M.M., et al., An integrated set-up for ex vivo characterisation of biaxial murine artery biomechanics under pulsatile conditions. Sci Rep, 2021. 11(1): p. 2671.

18. Santelices, L.C., et al., Experimental system for ex vivo measurement of murine aortic stiffness. Physiol Meas, 2007. 28(8): p. N39–49.

19. van der Laan, K.W.F., et al., Effect of rapid cooling, frozen storage, and thawing on the passive viscoelastic properties and structure of the rat aorta. Journal of Biomechanics, 2024. 171: p. 112190.

20. Caulk, A.W., J.D. Humphrey, and S.I. Murtada, Fundamental Roles of Axial Stretch in Isometric and Isobaric Evaluations of Vascular Contractility. J Biomech Eng, 2019. 141(3): p. 0310081–03100810.

21. Krasny, W., et al., Kinematics of collagen fibers in carotid arteries under tension-inflation loading. Journal of the Mechanical Behavior of Biomedical Materials, 2018. 77: p. 718–726.

22. Chen, H., et al., Biaxial deformation of collagen and elastin fibers in coronary adventitia. J Appl Physiol (1985), 2013. 115(11): p. 1683–93.

23. Roy, D., et al., A Literature Review of the Numerical Analysis of Abdominal Aortic Aneurysms Treated with Endovascular Stent Grafts. Computational and Mathematical Methods in Medicine, 2012. 2012: p. 820389.

24. Leloup, A.J.A., et al., Vascular smooth muscle cell contraction and relaxation in the isolated aorta: a critical regulator of large artery compliance. Physiol Rep, 2019. 7(4): p. e13934.

25. Neutel, C.H.G., et al., Empagliflozin decreases ageing-associated arterial stiffening and vascular fibrosis under normoglycemic conditions. Vascular Pharmacology, 2023. 152: p. 107212.

